# Correcting artifacts in ratiometric biosensor imaging; an improved approach for dividing noisy signals

**DOI:** 10.1101/2021.04.13.439673

**Authors:** Daniel J. Marston, Scott Slattery, Klaus M. Hahn, Denis Tsygankov

## Abstract

The accuracy of biosensor ratio imaging is limited by signal/noise. Signals can be weak when biosensor concentrations must be limited to avoid cell perturbation. This can be especially problematic in imaging of low volume regions, e.g., along the cell edge. The cell edge is an important imaging target in studies of cell motility. We show how the division of fluorescence intensities with low signal-to-noise at the cell edge creates specific artifacts due to background subtraction and division by small numbers, and that simply improving the accuracy of background subtraction cannot address these issues. We propose a new approach where, rather than simply subtracting background from the numerator and denominator, we subtract a noise correction factor (NCF) from the numerator only. This NCF can be derived from the analysis of noise distribution in the background near the cell edge or from ratio measurements in the cell regions where signal-to-noise is high. We test the performance of the method first by examining two noninteracting fluorophores distributed evenly in cells. This generated a uniform ratio that could provide a ground truth. We then analyzed actual protein activities reported by a single chain biosensor for the guanine exchange factor Asef, and a dual chain biosensor for the GTPase Cdc42. The reduction of edge artifacts revealed persistent Asef activity in a narrow band (∼640 nm wide) immediately adjacent to the cell edge. For Cdc42, the NCF method revealed an artefact that would have been obscured by traditional background subtraction approaches.

## Introduction

FRET biosensors are powerful, widely used tools for visualization and analysis of protein activities that include conformational changes, post-translational modification, and ligand interactions. Signaling proteins, like the Rho family GTPase and GEF studied here, function differently in their active and inactive conformations, so it is important to differentiate their overall distributions from the distribution of specific active conformations (Rossman, Der et al. 2005, Hall 2012). In Förster resonance energy transfer (FRET) biosensors, the conformational changes of the target protein typically modulate the separation or orientation of two fluorophores, a FRET donor and a FRET acceptor (Hochreiter, Garcia et al. 2015, Greenwald, Mehta et al. 2018, Terai, Imanishi et al. 2019). This produces a conformation-dependence of the FRET intensity (acceptor emission upon donor excitation). In ratiometric imaging, the FRET intensity is divided by the fluorescence intensity of the directly excited donor or acceptor fluorophore at each point in the cell. This ratio reflects the conformation of the target protein. Use of a ratio rather than simply measuring FRET intensity reduces artifacts produced by variations in cell volume or biosensor distribution (Kurokawa, Itoh et al. 2004, Pertz, Hodgson et al. 2006).

Processing FRET biosensor data to produce ratios involves a number of steps, each of which can affect the accuracy of the final result (e.g., shade correction, masking, subtraction of background fluorescence in each channel, correction for bleed-through between the channels, and correction for photobleaching (Machacek, Hodgson et al. 2009, Hodgson, Shen et al. 2010)). Here we focus on artifacts produced when noise and background subtraction exert strong effects on the final ratio. Errors introduced by oft-used procedures can create artefactual, apparent gradients in the ratio near the cell edge, a region particularly important when studying cell morphodynamics. When relatively flat cells are used to study motility, important actin dynamics occur in a thin region within 2-3 microns of the edge (Pertz, Hodgson et al. 2006, Machacek, Hodgson et al. 2009, Marston, Vilela et al. 2020).The small volume of this region decreases signal/noise and increases the magnitude of background subtraction errors relative to the real signal. Importantly these effects increase as we move towards the cell edge and volume decreases. We show here how and why this region is prone to artefacts, and propose a correction method, the noise correction factor (NCF), that eliminates the need for background subtraction. The method can be applied in many types of ratio imaging where background subtraction can be problematic.

The paper begins with a simplified model of the problem, based on considering a hypothetical cell with thickness gradually decreasing as we move closer to the edge. This model is used to illustrate issues with background subtraction and noise that generate artefactual gradients in ratios, and to explain the rationale behind the NCF method. We demonstrate the artefactual gradients in real cells and look more closely at the role that background subtraction plays in such artefacts. Using an “inert biosensor” that would be expected to produce a uniform ratio throughout the cell, we demonstrate why improving the accuracy of background subtraction actually amplifies the artefactual gradient associated with division by low signal-to-noise values. Finally, we provide mathematical derivations that justify the use of NCF to produce an artifact-free ratio. These theoretical considerations are tested by applying the NCF method to two additional biosensors: a single-chain biosensor for Asef, and a dual-chain biosensor for the GTPase Cdc42. For the Asef biosensor, by eliminating the noise-related artifact we eliminated spurious activity at the cell edge, revealing real GEF activation in a 2-pixel wide band along the periphery. For the GTPase biosensor, improved visualization of near edge regions revealed problematic areas that needed to be excluded from ratio analysis.

## Results

### A hypothetical cell model illustrating effects of noise on ratio values

Here we consider a simple cell model to establish the new approach; it will be applied to real cells in the following sections. FRET is defined as the fluorescence intensity emitted from the acceptor upon donor excitation, excluding any contribution from background, spectral bleedthrough or other artifacts. Emission from the “FRET channel”, on the other hand, is the intensity measured when the microscope is configured to monitor donor excitation and acceptor emission, including contributions from bleedthrough, background etc. Analogously, monitoring the “Donor channel” refers to quantifying light actually collected when monitoring donor excitation and donor emission, including artefactual contributions. Let us consider a hypothetical cell in which the thickness gradually increased with the distance from the edge, starting with a value of zero thickness at the edge. Let’s assume that the protein activity, as reflected by a FRET biosensor, is perfectly uniform throughout the volume of the cell. Ideally, the intensity of the signals in the FRET and donor channels will increase linearly with the distance from the edge, *x*, as *S*_*F*_*x* and *S*_*D*_*x* respectively, due to the increasing thickness of the cell. By taking the ratio of the signals, we get the constant value 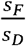, which tells us that the activity is uniform as expected. In reality, each signal is shifted by a background fluorescence: *S*_*F*_*x* + backgraound and *S*_*D*_*x +* backgraound. Thus, inaccurate subtraction of the background leads to the ratio 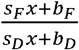, which is a function of *x* instead of a constant 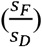. The more accurately we subtract the background, the more accurately we can determine the ratio. Let’s assume that we found a way to subtract the background perfectly. Now we have another problem: each channel has some noise in the signal so that the ratio after background subtraction is 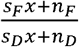. Far enough from the edge, the signal dominates the noise, (*S*_*F*_*x* ≫ *n*_*F*_ and *S*_*D*_*x* ≫ *n*_*D*_), so we get an accurate estimate of 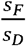. However, near the very edge, as *x* → 0, the ratio is heavily affected by noise, producing large variance of the ratio values. The calculated ratios will tend to produce more large positive values when noise starts to dominate the signal (effects of noise that decrease the correct ratio value can only approach zero, whereas those that increase the ratio can become arbitrarily high). Importantly, this trend towards increasing values will have a spatial component, making the ratio appear to become higher as we approach the edge. The more high-intensity pixels are produced by this effect, the higher will be the apparent, artefactual increase in activity at the edge, which be mistaken for a real protein activity gradient.

In this example, we can resolve these issues using the fact that 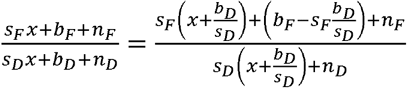 (**Supplementary Information**). Instead of subtracting background values from each channel, we can subtract one *correction factor*, 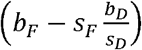, from the FRET channel only, producing the ratio 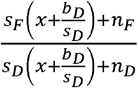, which is a better estimate of 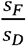, because even as *x* → 0, 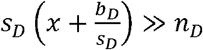 and we never divide by noise.

In the following sections, we explore the applicability of this idea to actual biosensor data. The correction factor will be referred to as the noise correction factor (NCF).

### Effects of noise examined in real cells

To illustrate the effect of noise in wide field imaging of real cells, we examined Cos7 cells expressing two fluorescent proteins often used for FRET, mCerulean and CyPet. Although biosensor activity is frequently reported as the ratio of FRET intensity to the intensity of the directly excited donor fluorophore, other ratios are also used (e.g., FRET divided by a directly excited acceptor or a volume indicator). To keep our description more general, we refer to the numerator as image1 and the denominator as image2. When background subtraction is explicitly included, the ratio is:

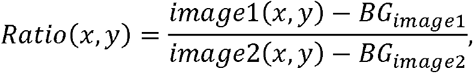

where (*x, y*) is the position of a pixel in the image.

**Figure 1A** shows the nonuniform intensity distribution in the FRET channel for two neighboring cells, containing linked fluorophores that should FRET but not reflect protein activity. **Figure 1B** shows the fluorescence ratio after background subtraction for the same cells, a much more uniform image. For the background of each image, we determined the average intensity of a region away from the cell, which we will refer to as a *distant background* **(green box in Figure 1A)**. To more clearly show variations in ratio values, we set the pseudocolor scale by assigning the region outside the cells to equal the mean of the ratios inside (≈ 0.7). A line-scan across the cells (black line in Figure 1B) showed that some pixels at the cell edges had intensity values several-fold higher than those within the cell (arrows in **Figure 1C**), as predicted. **Figure 1D,E** show that such pixels create a statistical bias in the average intensity within the cell, which varies with the distance from the edge,*d*. Using the relative range of the mean intensity, (max_*d*_ ⟨*I*⟩ − min_*d*_⟨*I*⟩)/ min_*d*_⟨*I*⟩, as the measure of the deviation from the expected constant mean, we find that the left and right cells are biased at their edges by 4.7% and 5.7%, respectively.

**Figure 1.**
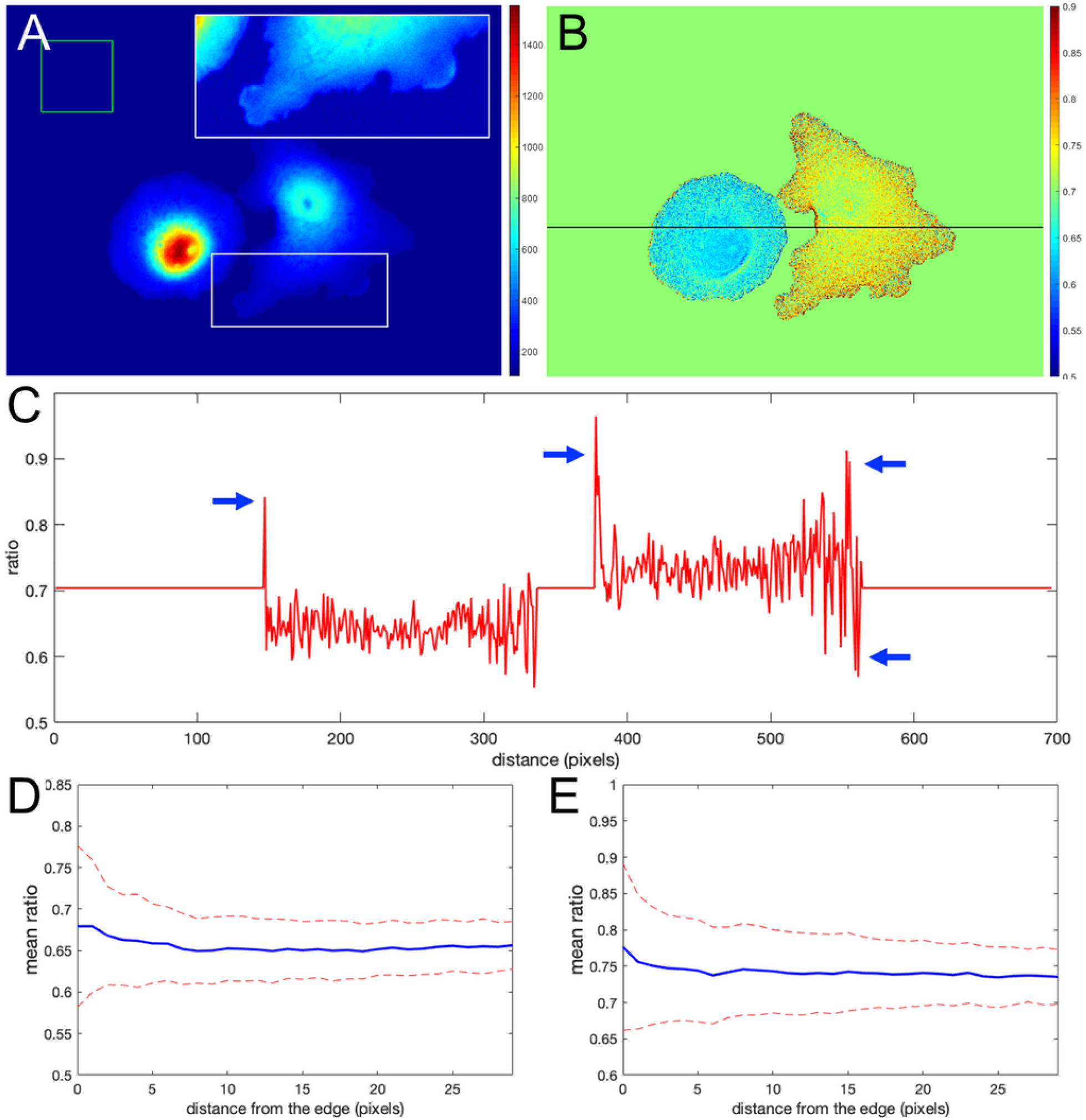
Ratio calculation using the mean background subtraction method. **A**. A colormap of the FRET channel (mCerulean emission upon excitation of CyPet) after background subtraction, showing noneven distribution near the cell edge. The insert shows the signal from the white box in log scale for better contrast at the edge. The green box indicates the area of the image used to determine the mean and standard deviation of the background noise. **B**. The resulting ratio signal with the background set to the mean ratio signal in the cells for a closer look at the non-uniform distribution. **C**. The ratio profile from B along the black line. Blue arrows point to the artifact – at the edge very large deviations of the ratio from the expected constant mean level in the cells. **D, E**. Mean ratio signal as a function of the distance toward the cell center from the edge for the left and right cells in B. Blue curves indicate the mean. Red dashed curves indicate the mean plus/minus one standard deviation.

### The effects of background subtraction

Like noise, incorrect background subtraction can also generate artefactual gradients at thin parts of the cell. Importantly, the method we will propose here does not require direct background subtraction, so these artefacts will be eliminated. In this section we show why it is worth eliminating the need for background subtraction; and highlight potential artifacts for those who do use it.

Incorrect background subtraction artificially elevates or reduces ratios, depending on the relative extent of the error it introduces in the numerator and denominator. As the signal decreases towards the edge, such background errors contribute more to the ratio values calculated, producing an artefactual gradient of activity (see Figure 2 for an extreme case of no background subtraction at all). The gradient can increase or decrease towards the cell edge, depending on the relative magnitude of errors in the denominator and numerator. To put it mathematically, as the signals and get smaller near the edge, the errors in the background subtraction make that ratio _____________________ approach a number _____, which has no biological meaning.

**Figure 2.**
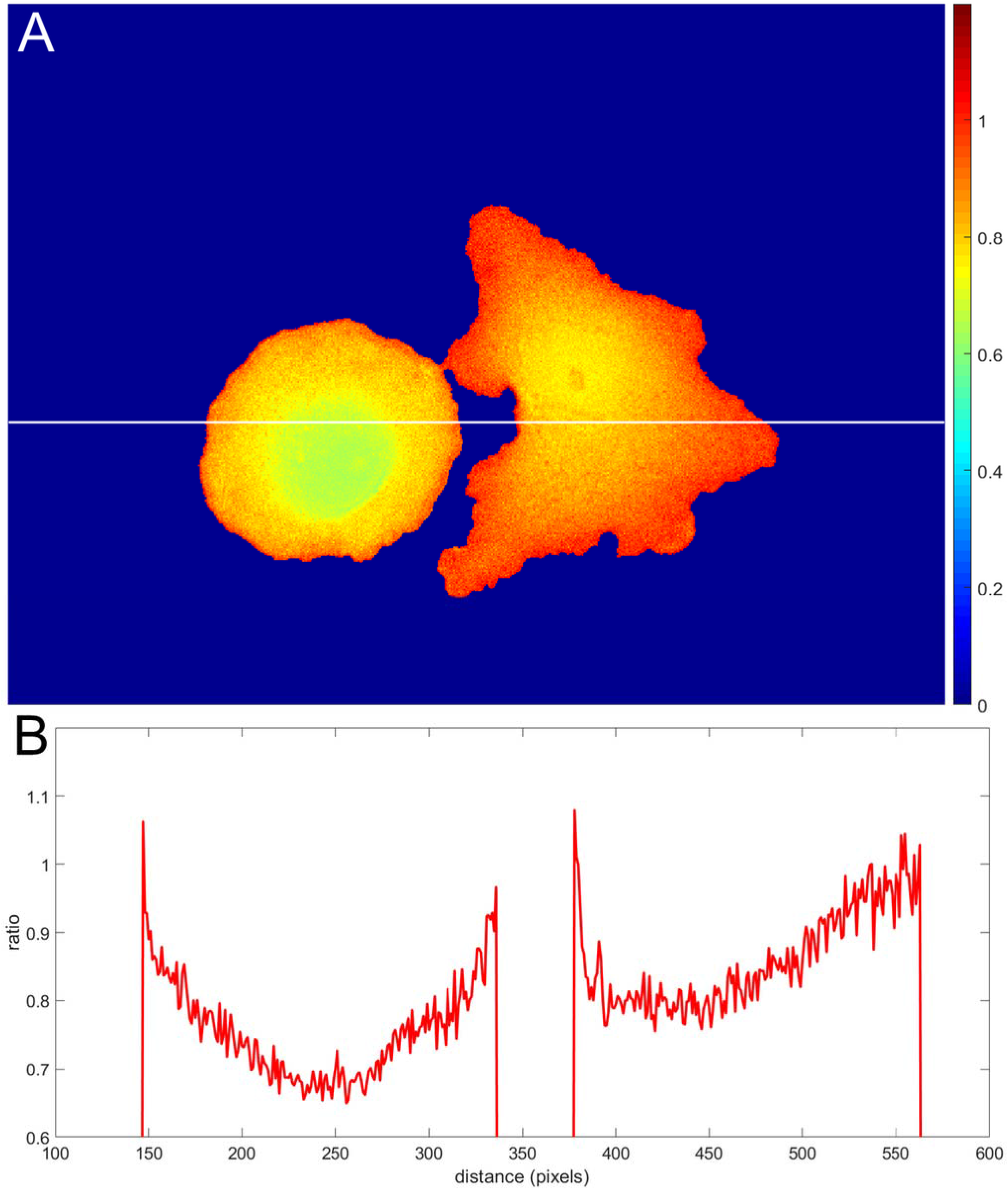
Illustration of an extreme case of an error in background subtraction. **A**. The ratio _____without background subtraction at all **B**. The ratio profile from A along the white line.

Determining the correct background to apply to each pixel in the cell can be challenging. In wide field imaging for example, parts of the cell outside the plane of focus generate out-of-focus light that is unevenly distributed across the in-focus pixels. Some of this light appears outside the cell, so averaging regions outside the cell to determine the background can be problematic. When averaging a region far from the cell edge, the background will be too low. To illustrate this effect, we compared the ratios obtained in Figure 1 with those obtained by setting the backgrounds (and) halfway between the distant background value used for Figure 1, and the value used to define the cell mask, i.e., the value right at the cell edge. **Figure 3A,B** shows that the resulting bias becomes worse, giving a deviation of 4.2% and 10.6% from the flat level, on average, and the variance of the ratio values at the edge is significantly increased. This was because using an increased background value decreased the intensity after background subtraction, making it smaller relative to noise.

**Figure 3.**
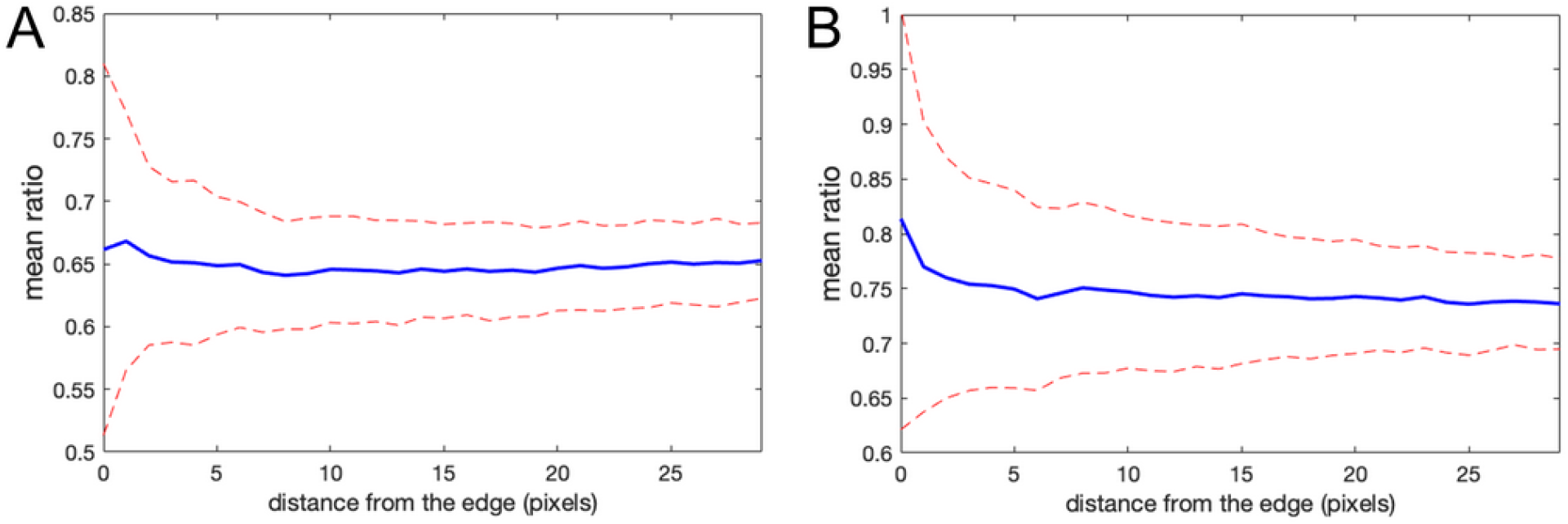
Mean ratio signal as a function of the distance from the edge. Blue curves indicate the mean. Red dashed curves indicate the mean plus/minus one standard deviation. **A,B**. The result for the left and right cells in Figure 1 based on the subtraction of the values halfway between the distant background value and the value used to define the cell mask. Here the deviations from the expected constant mean are 4.2% and 10.6%, respectively.

We tested whether this problem could be overcome by subtracting background values obtained near the edge of the cell, taking into account the fact that background near the edge varies along the periphery of the cell. We applied *non-uniform background subtraction* by subtracting and that depend on the position in the image. To capture the spatial variation of the background, the background intensity was measured along the cell edge right outside the cell, and the resulting values were applied to nearby regions within the cell, so that subtracted values more properly reflected the local background. **Figure 4** illustrates the approach.

**Figure 4.**
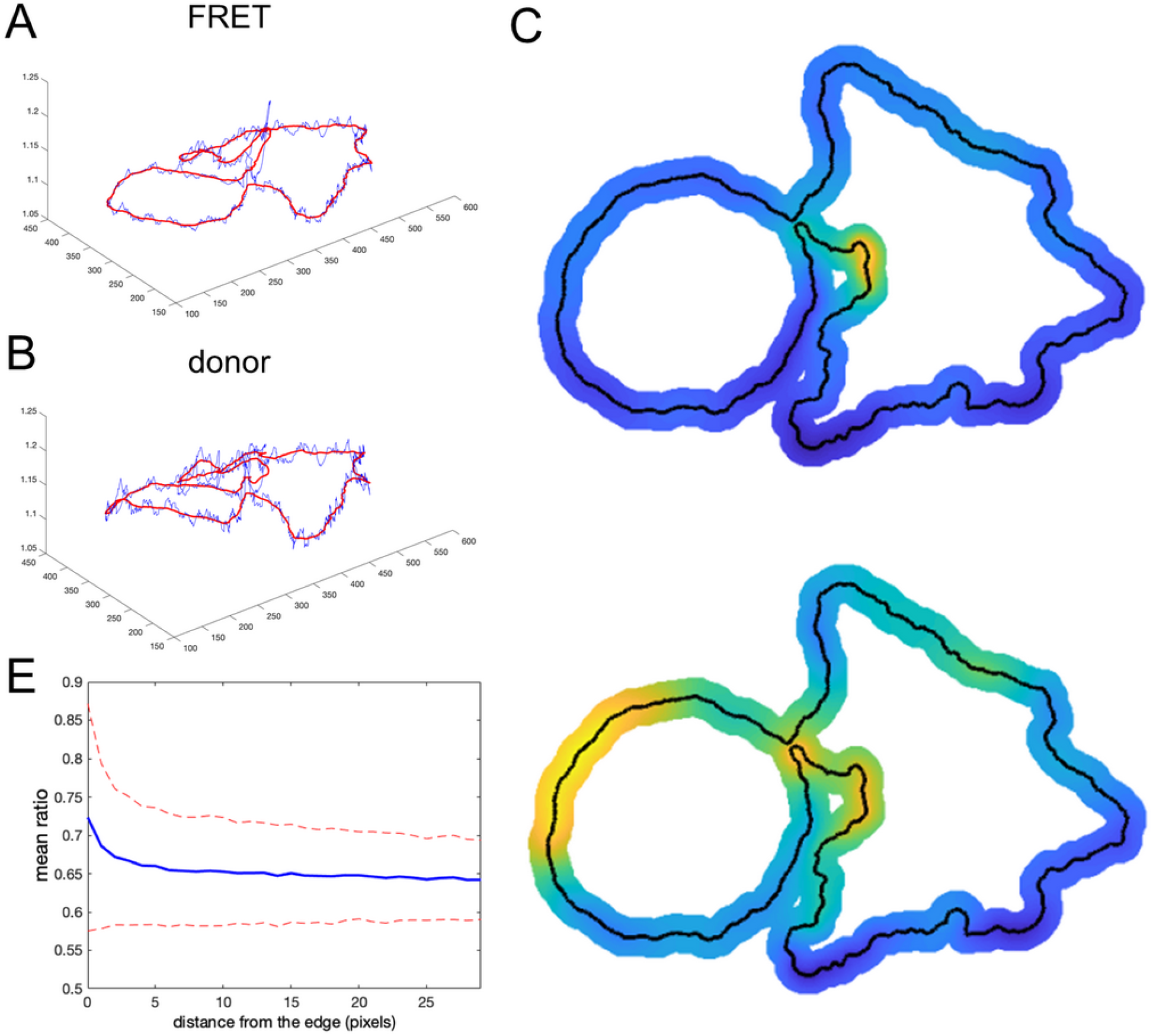
Non-uniform background subtraction. **A,B**. The fluorescence intensity in the FRET and donor channels outside the cell and near the edge (i.e., the local background along the cell periphery) is shown in blue. Smoothing these using a Gaussian filter generated the curve shown in red. **C,D**. Spatial interpolation using the smoothed signal along the cell edge as the background for determining ratio values near the edge. **E**. Mean ratio signal as a function of the distance toward the cell center from the edge, showing 12.7% deviation from the expected constant mean. This is due to the same edge artifacts shown in Figure 1, but further exaggerated by small values of the ratio denominator.

First, we find the intensity at each pixel adjacent to the cell boundary for each channel (blue curves in **Figure 4A,B**). This measurement reflects two contributions: 1) local pixel-to-pixel variation due to the intrinsic noise of the signal and 2) a larger-scale variation in the background at different regions around the cell. To estimate background fluorescence along the whole cell edge, we need to use this larger-scale trend in the intensity variation (red curves in Figure 4A,B). It is found by applying the Gaussian filter(Davies 2005) to smooth out the noise, _ ____ ___, where and are numerical indexes of the points along the edge and is a parameter representing the extent of the smoothing window. Next, we find the interpolated values inside the cell as ______, where is the distance from a pixel *i* to the point (*x, y*) and *m* is the parameter that controls how deep the local background values at the cell edge are extended inside the cell before the peripheral variation is smoothly connected across the cell. Such interpolation produces a meaningful estimation of the background intensity distribution near the curved edge. Interpolated distribution in the middle of the cell may not be any more accurate than a global value of the background obtained from a distant region. The result of this operation is shown in **Figure 4C,D** for the FRET and donor channels, respectively (only the near edge regions are shown). Finally, we find the ratio after non-uniform background subtraction as:

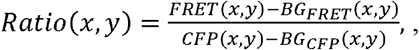

Designed this way, our non-uniform background should provide a particularly accurate estimate of the actual background signal around the cell near its edge. However, the problem with the division by small numbers (weak donor signal at the very edge) is not resolved. The artifact is actually stronger because the subtracted values are closer to the fluorescent signal on the cell edge. Indeed, **Figure 4E** shows the mean intensity change in the cell with the distance from the edge. Our metric of the deviation from the expected flat distribution (max_*d*_ ⟨*I*⟩ − min_*d*_⟨*I*⟩)/ min_*d*_⟨*I*⟩, gives 12.7%, which is about twofold more than the result of the distant background subtraction method (compare with **Figure 1D,E**).

### Use of a noise correction factor; identification and correction of artifacts without using direct background subtraction

The previous section shows that improving background subtraction may not be sufficient to avoid misleading ratiometric artifacts at the edge of the cell. Let us therefore explore an alternative way to calculate the ratio of image 1 and image 2. We start with a theoretical estimation and then test the predictions with live-cell imaging.

Let’s start with the situation considered above, where protein activity is constant throughout the cell, so that signal variation across the cell results from non-uniform cell thickness. We aim to determine the ratio between background- and noise-free florescent signals *S*_1_(*x, y*) and *S*_2_(*x, y*), which must be proportional to each other in this case. The ratio of the two images can be written as

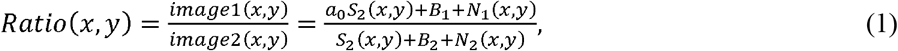

where *a*_0_ is a coefficient of proportionality and *B*_1_,*N*_1_ (*x, y*),*B*_2_,*N*_2_ (*x, y*) are the background levels and the noise in image 1 and image 2, respectively.

Background subtraction from each image (e.g., using mean background subtraction, MBS) reports:

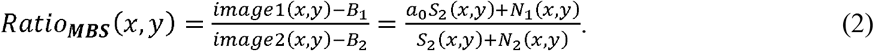

Using a simple algebraic rearrangement of terms in this equation (1), we find (see **Supplementary Information**)

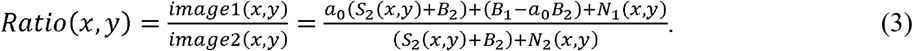

*Now we can show that subtracting a specific constant in the numerator – the noise correction factor (NCF) – eliminates the need for background subtraction in the denominator*. Subtracting *NCF* = *B*_1_ − *a*_0_*B*_2_ produces:

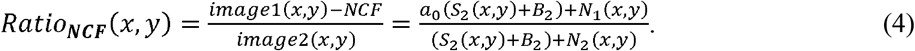

In any part of the cell away from the edge, where *S*_2_(*x, y*) ≫ *B*_2_ ≫ *N*_2_(*x, y*), both the MBS and NCF methods give the same correct (flat on average) result:

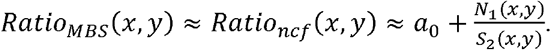

However, in thin regions near the edge, where *S*_2_(*x, y*) → 0, the MBS method generates the artifact

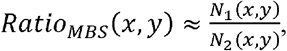

while the NCF method still gives the correct (flat on average) answer

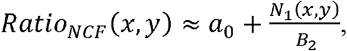

When we use the correction factor, the artefact is not present because we did not subtract the background from the denominator and *B*_2_ ≫ *N*_2_(*x, y*). For the same reason, we can apply the correction factor approach to the whole image, including the region around the cells (with no biosensor signal). For these regions, we get:

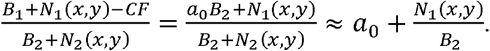

So far, we considered an ideal (biologically uninteresting) situation where the FRET signal is strictly proportional to the donor signal. Now, let’s consider the next order of approximation, where biosensor activity deviates from the basal level so that the pure (background- and noise-free) part of the signal in image 1 is *S*_1_(*x, y*) = [*a*_0_ + *a*_1_(*x, y*)] *S*_2_(*x, y*). As we will illustrate below, when we discuss imaging of a GEF biosensor, this is a good approximation for biosensors that are based on a single protein chain which contains both FRET fluorophores. Subtracting the correction factor *NCF* = *B*_1_ − *a*_0_*B*_2_ from image 1 yields

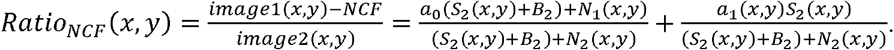

In cell regions where the donor signal is strong *S*_2_(*x, y*) ≫ *B*_2_ ≫ *N*_2_(*x, y*), we again get the agreement between the MBS and NCF methods:

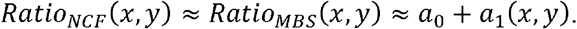

In the background of the image, we still get

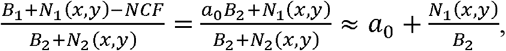

while at the very edge of the cell, where *S*_2_(*x, y*) → 0, but *B*_2_ ≫ *N*_2_(*x, y*), the corrected ratio transitions between *a*_0_ and *a*_0_+ *a*_1_(*x, y*) as

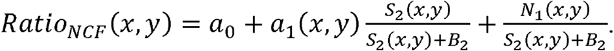

These results suggest that in cell areas where the MBS method does not generate noise-related artifacts, the noise correction factor gives the same ratio values. However, unlike the MBS method, the NCF method allows us to evaluate the ratio all the way to the cell edge. In addition, the NCF method allows us to visualize background noise outside the cell (Figure 5, 6). This is useful because it enables comparison of background noise outside the cell with the FRET ratio inside the cell. With the MBS method, this is not possible because outside the cell we have one noise divided by another noise. Although the NCF method may underestimate the values at the very edge of the cell, these values would still be appreciably higher than the zero activity FRET ratio *a*_0._ FRET signal measured when there is no protein activity reported by the biosensor could result from the FRET of the biosensor in its ‘off state’ (e.g., for single cell biosensors where both fluorophores are always held in proximity within the biosensor) or from some systematic shift in the image acquisition (e.g., an uncorrected camera signal). The practical usefulness of this method is to visualize the ratio signal at the cell edge and other noisy portions of the cell, free from artifacts associated with division by a very weak donor signal.

**Figure 5.**
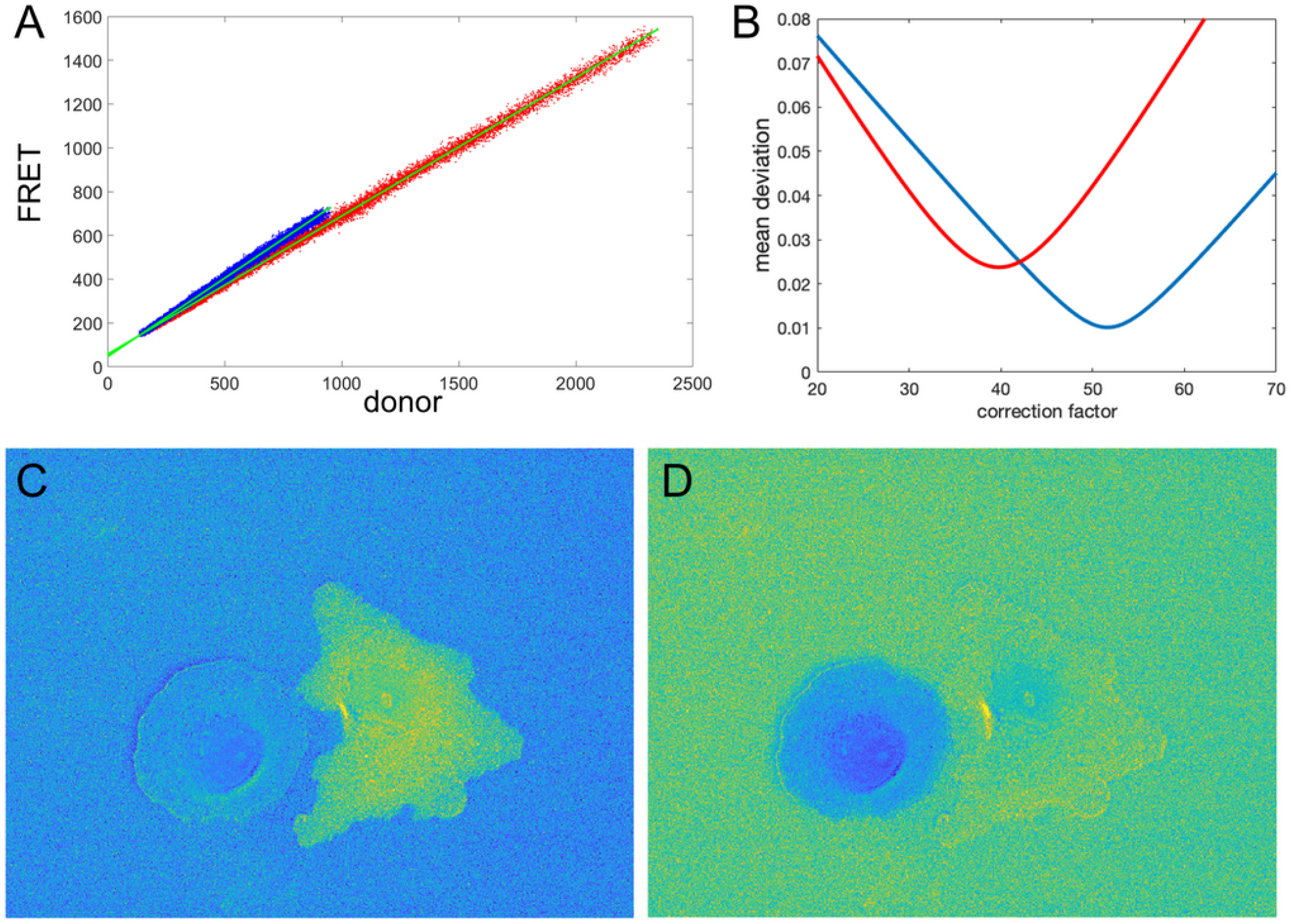
Illustration of the correction factor method. **A**. Intensity in the FRET vs. donor channels for each pixel. Red and blue colors indicate pixels in the left and right cells, respectively. The green lines indicate linear fits to the data points. **B**. The deviation function (Equation 5) for a range of correction factor values. Red and blue curves are the functions calculated separately for the left and right cells in the image. The optimal correction factors are defined as the positions of the minima of these functions. **C,D**. The ratio images resulted from the subtraction of the correction factors. In C, the signal in the left cell flattens with the background level as predicted by the mathematical analysis. Similarly, in D, the right cell flattens with the background. The flattening is so efficient that the noise fluctuations become visible.

**Figure 6.**
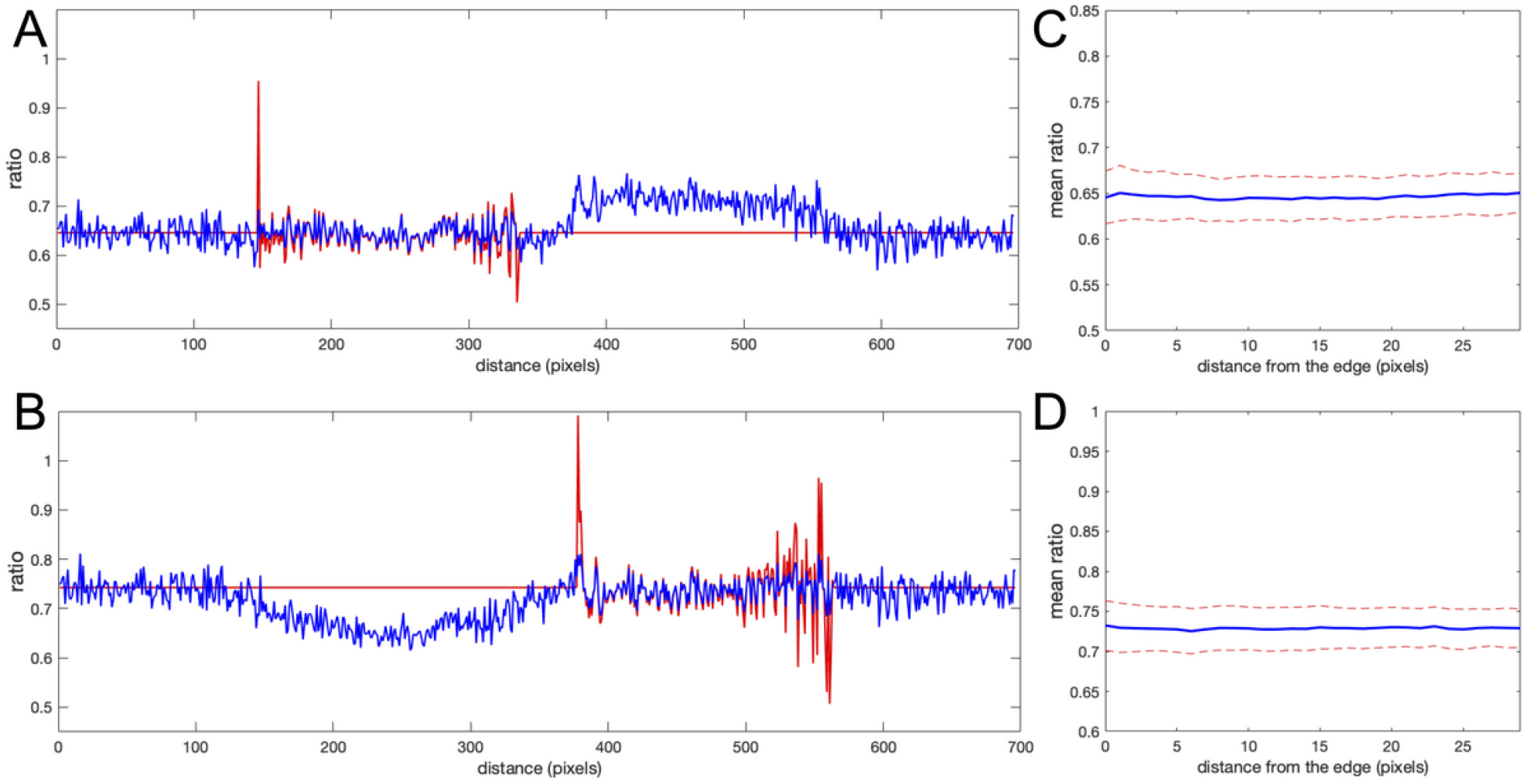
Comparison of correction using the noise correction factor (NCF) versus mean background subtraction. **A,B**. The ratio profiles from the images in Figure 5C,D along the line at the same location as in Figure 1. Red and blue colors indicate the ratios obtained with the mean background subtraction and NCF methods, respectively. Both results overlap inside the cells away from the edge (left cell in A and right cell in B). Near the cell edges, the shape of the profile is still consistent between the methods, but the artifacts are not present in the result of the correction factor method. **C,D**. Mean ratio signal as a function of the distance toward the cell center from the edge after applying the correction factor (C for the left cell and D for the right cell), showing the expected flat level with only 1.2% and 0.9% deviation from a perfectly constant line. The variance of the fluctuations is also flat all the way to the cell edge, in contrast to the background subtraction approaches (see Figure 3 and Figure 4D).

### Determining the proper noise correction factor in practical applications

The mathematical results in the previous section illustrated the benefits of using the NCF method. Here we show two alternative ways to find the right value of NCF for practical applications. In some cases, there will be appreciable “background FRET”, measurable intensity in the FRET channel even when there is no protein activity. This could occur, for example, with single chain biosensors that contain two fluorophores held near each other in the biosensors ‘off’ state (Pertz, Hodgson et al. 2006, Marston, Vilela et al. 2020). In such cases, we can take advantage of the fact that the correct NCF value makes *Ratio*_*NCF*_(*x, y*) ≈ *Ratio*_*MBS*_(*x, y*) in the cell regions away from the edge. It is easy to check for strong background FRET by plotting the intensity of image 1 vs. image 2 for each pixel. In such cases, the NCF can be found as the value that minimizes the deviation function

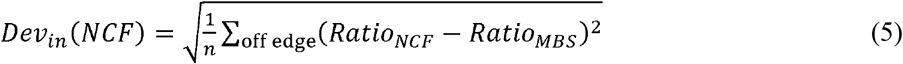

where *n* is the number of pixels in the considered cell region away from the edge.

Another way to determine the NCF is to take advantage of the fact that the correct NCF value makes the background level near the edge outside the cell flat: 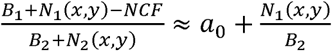. Thus, the NCF can be found as the value that minimizes the deviation function

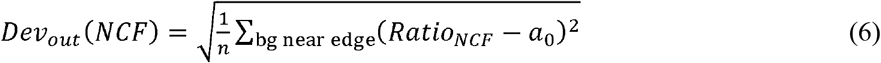

where *n* is the number of pixels in the considered background region. Since *a*_0_ represents the zero protein activity “background ratio”, we define it as the mean value of the ratio calculated with the MBS method inside the cell. This approach is preferable for biosensors where we cannot expect strong proportionality between background- and noise-free florescent signals *S*_1_(*x, y*) and *S*_2_(*x, y*), e.g., for biosensors consisting of two chains, with one fluorophore on each chain (Machacek, Hodgson et al. 2009, Marston, Vilela et al. 2020).

Let us check how this optimization approach works for our example from the previous sections (two fluorophores with constant FRET throughout the cell). We first verify the proportionality between the signals in image 1 and image 2. **Figure 5A** shows FRET vs. donor signals for each pixel in the image with the expected strong proportionality. The linear fit gives the coefficients *a*_0_ = 0.63 and *a*_0_ = 0.72 for the left and right cells in the image, respectively. Knowing the background values, we can estimate the theoretical value (*B*_*F* −_ *a*_0_*B*_*D*_) of the noise correction factors as *NCF* = 53.4 and *NCF* = 43.2. Next, using morphological erosion (Soille, P. 2004) of the cell masks by 10 pixels, we determine the deviation values of the function (5), as shown in **Figure 5B** for each of the cells. The smallest deviation values are achieved when *NCF* = 51.6, and *NCF* =39.8, which are close but not exactly equal to our estimation based on the fit of the FRET vs. donor plot. **Figure 5C,D** shows the resulting ratio images, and **Figure 6A,B** shows the corresponding line scans across the image.

Clearly, the MBS and NCF methods give the same intensity profile, except that the NCF method does not show noise-related artifacts, in agreement with the theoretical prediction. For and we found the smallest value of the flatness metric,, when the mean intensity became close to a constant value over a range of distances from the edge (**Figure 6C,D**), as expected for this data. For these correction factors, the deviation from the constant became 1.2% and 0.9% for the left and right cells in the image, respectively. This is a fivefold improvement over the MBS method (see Figure 3A,B). All the values of NCF for each cell found through different means above are consistent with each other, which supports the mathematical rationale behind the NCF method.

### Application of the correction factor method to a single-chain biosensor

We next tested the theoretical predictions of the previous sections using our single-chain biosensor for the guanine exchange factor (GEF) Asef (Marston, Vilela et al. 2020). In this biosensor, a pair of fluorophores, mCerulean 3 and YPet, connected by a flexible linker, are inserted into a flexible hinge region between the active site and autoinhibitory domain (AID). Activation of the protein causes the AID to be displaced and the donor/FRET ratio to increase. The biosensor was imaged in moving fibroblasts constitutively expressing the biosensor. **Figure 7A** shows the plot of FRET vs CFP values for each pixel of a biosensor image. Clearly, there is a strong linear trend due to the zero-activity background ratio of the connected fluorophores, with a number of pixels deviating from the line due to biological activation,, as can be seen in the zoomed region of the plot (**Figure 7B**). Therefore, our theoretical representation of this FRET vs. CFP relationship as ____ is a good approximation. Now, **Figure 7C** shows the ratio of the two channels obtained with the MBS method. Close visual inspection of the pixels near the edge of the segmented cell (see **Figure 7D,E**) indicates that there are many very bright, somewhat irregularly distributed pixels along the edge. These pixels may well have been generated by the artifact that we investigated in the first sections.

**Figure 7.**
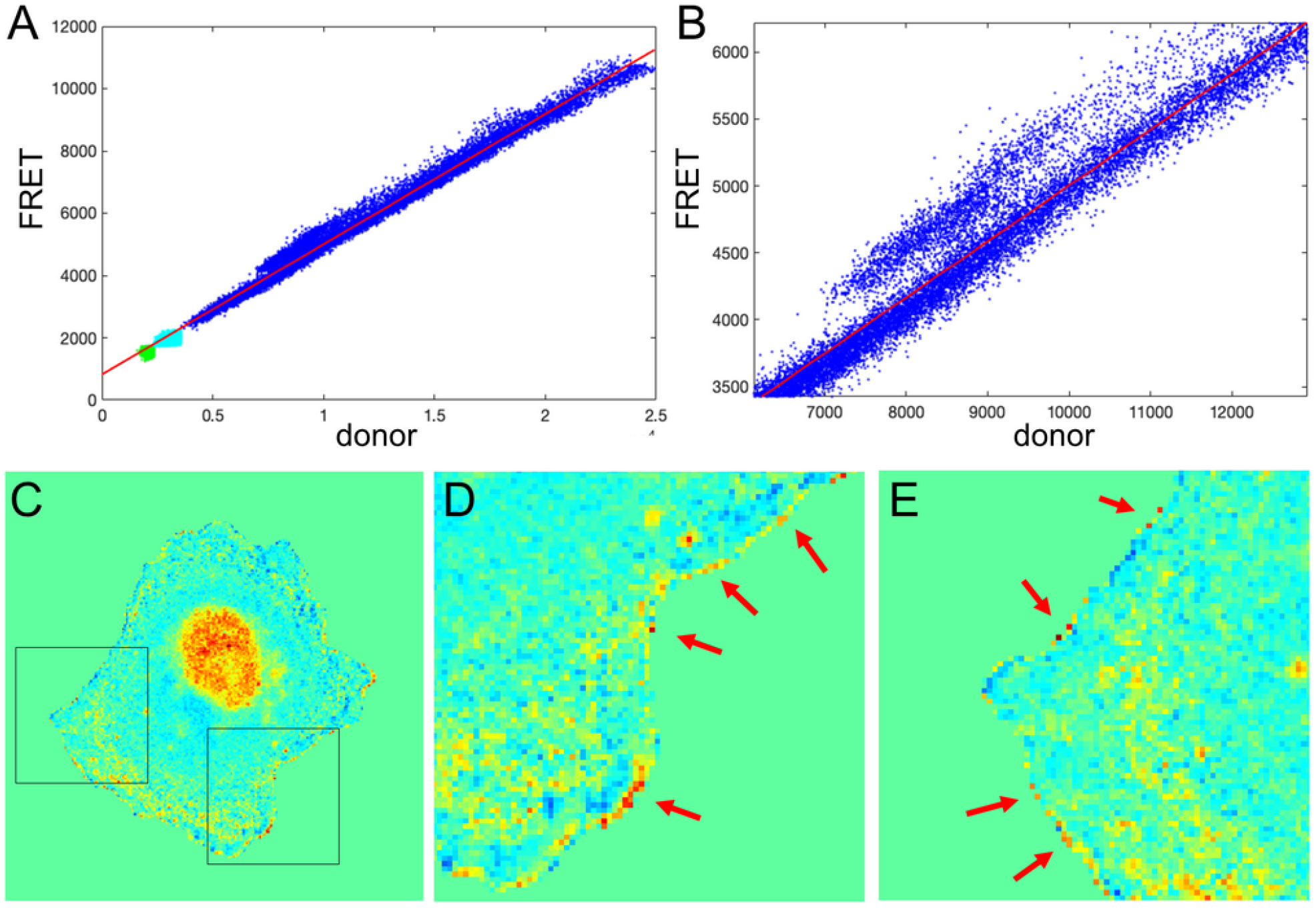
Application of the mean background subtraction method to Asef biosensor data. **A**. Uncorrected FRET vs. donor for each pixel. Blue indicates pixels with intensities above the mask threshold (i.e., inside the cell), green indicates pixels in the background with intensities below the mean plus three standard deviations of the noise in the box, and cyan indicates pixels in the background near the cell edge (with intensities between the threshold and mean + 3 std of the background noise) **B**. A zoomed-in region of the image in A, confirming our theoretical assumption that there is strong basal activity of the GEF biosensor. **C**. Ratio image resulting from the mean background subtraction method. Here we set the background level to the mean value of the rati inside the cell (as in Figure 2B). **D,E**. Two different zoomed-in regions from the image in C (black boxes). Red arrows point to high ratio pixels on the edge of the cell. With MBS method, these pixels appear similar to the noise artifact seen Figure 1C, so NCF is needed to analyze these pixels properly.

How would we know if this is real biosensor activity or an artifact stemming from ratio calculation? The correction factor approach can help to answer this question. First, we use the deviation metric (Equation 5) to find the noise correction factor. As described in the previous section, by minimizing the deviation we find the correct NCF, for which the ratio values away from the edge (where signal to noise is high) match the values of the MBS method (see **Figure 8A,B**). Now, we can calculate the ratio values across the whole image (inside and outside of the cell). With the NCF method, the values of the pixels at th very edge are significantly lower than the values of the same pixels in the MBS method. We can consider these values to be free of noise-related artifacts. However, they are still clearly elevated relative to th background level. Thus, the NCF ratio image indicates that there is indeed a narrow band of high activity on the edge of the cell (see **Figure 8C,D**), which is not a processing artifact but a true biological activity. For a better quantitative measurement of this effect, we plot the confidence regions for mean intensity values of the contour pixels on the cell edge and up to 2 pixels away from the edge toward the cell center and towards the cell background (**Figure 8E**). This plot shows that the band of activity on the cell edge is just 2 pixels (0.64 um) wide, and it stands out over the mean intensities on each side of the band with *statistical significance*.

**Figure 8.**
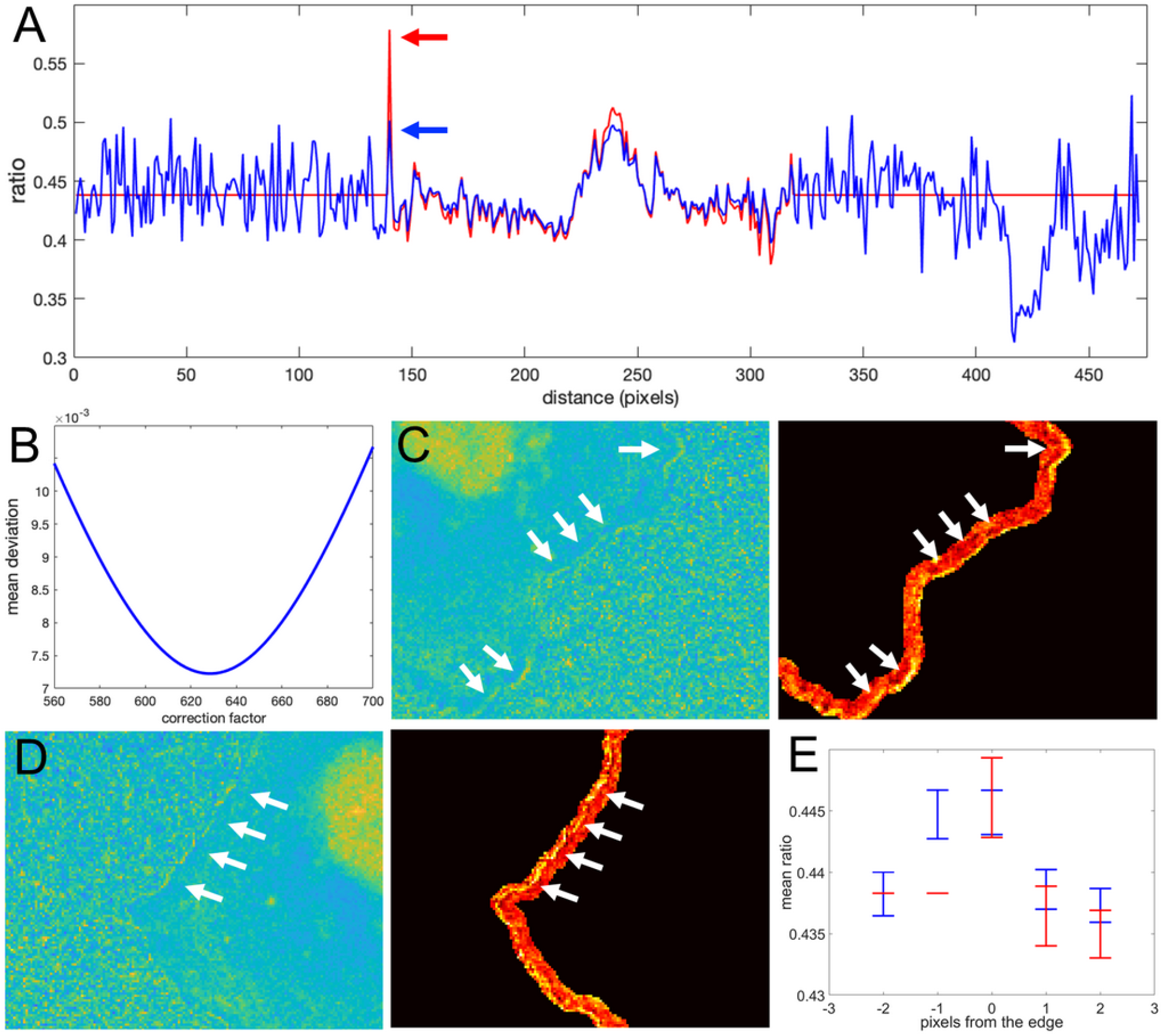
Application of the correction factor method to Asef biosensor data. **A**. The ratio values along a line in the ratio image resulting from the mean background subtraction method (red) and from the NCF method (blue). The red arrow indicates a pixel where the MBS method gives an artefactually high ratio value (noise-related artifact). The NCF method gives a value (blue arrow) that is within the distribution of ratio values seen throughout the cell (free of the noise related artifact). **B**. The deviation function (Equation 5) for a range of NCF values. The correction factor giving the minimum deviation is used. **C,D**. Two different zoomed-in regions of the ratio image calculated with the NCF method. Each zoomed-in region is presented in two colormaps. For better contrast, the panels on the right also exclude all regions from the view except for the near edge region. White arrows point to a narrow band of bright pixels at the very edge of the cell. **E**. Statistical measurement of the activity distribution (mean plus/minus the standard error) for pixels along the contours at different distances from the cell edge. Blue and red color represents the result of the NCF and MBS methods, respectively. On average, a two-pixel band is elevated over the noise seen beside the band, either inside or outside the cell.

### Application of the correction factor method to a dual-chain biosensor

Our theoretical considerations relied on the assumption that there is a strong component in the FRET signal that is proportional to the donor signal. As we showed in the previous section, this assumption is accurate for the single chain Asef biosensor and should be accurate in general for single chain FRET biosensors due to their design. For dual-chain biosensors using intermolecular FRET, we cannot expect the same level of correlation because the donor and acceptor are not physically linked and may or may not be distributed similarly in the cell. However, we can still find the NCF based on the flattening of the background level near the cell edge outside the cell, using the deviation function defined by Equation 6. This background level can be used as a baseline with which to compare ratio values inside the cell. Notice that in all our previous examples, the mean level of the noise outside the cell becomes flat on average and levels up to the mean ratio inside the cell after subtracting the NCF (See Figure 6A,B and 8A). This makes sense, because noise outside the cell should be flat on average regardless of the type of biosensor we use. Based on this minimization routine, we can expect that the proper NCF will suppress noise-related edge artifacts in ratiometric analysis of dual-chain biosensors. To investigate this, we applied it to the dual chain Cdc42 biosensor designed to be imaged simultaneously with the Asef biosensor. In this Cdc42 biosensor (Marston, Vilela et al. 2020), LSSmOrange is attached to Cdc42, and mCherry is attached to the CRIB domain of WASP. When Cdc42 is activated the WASP fragment binds Cdc42, and mCherry (FRET) emission can be detected upon LSSmOrange (donor) excitation. This biosensor could be used together with the Asef biosensor because the donor proteins in the two biosensors can be excited with the same wavelength, but they have substantially different emission maxima. Because this was a dual chain biosensor, we studied FRET and donor signals after correction for spectral bleedthrough. Ratios were then calculated using the MBS and NCF methods.

To define the region near the cell edge outside the cell, we used morphological dilation by 50 pixels. The minimum of the deviation function was achieved at *NCF* = 550.6 The resulting ratio images from the two methods are shown in **Figure 9A-C**. Perhaps surprisingly, the intensity profile inside the cell away from the edge was very consistent between the methods (**Figure 9D**), although here we did not use such criteria for finding NCF. Actually, if we do use the minimization based on the deviation function inside the cell, *Dev*_*in*_(*NCF*) (see Equation 5), we get the correction factor *NCF* = 538, which is close to the value we obtain with the minimization of *Dev*_*out*_(*NCF*). This result, confirming the agreement of the two types of optimization, further justifies our rationale for using flattening of the background level as a way to find the proper value of the NCF.

**Figure 9.**
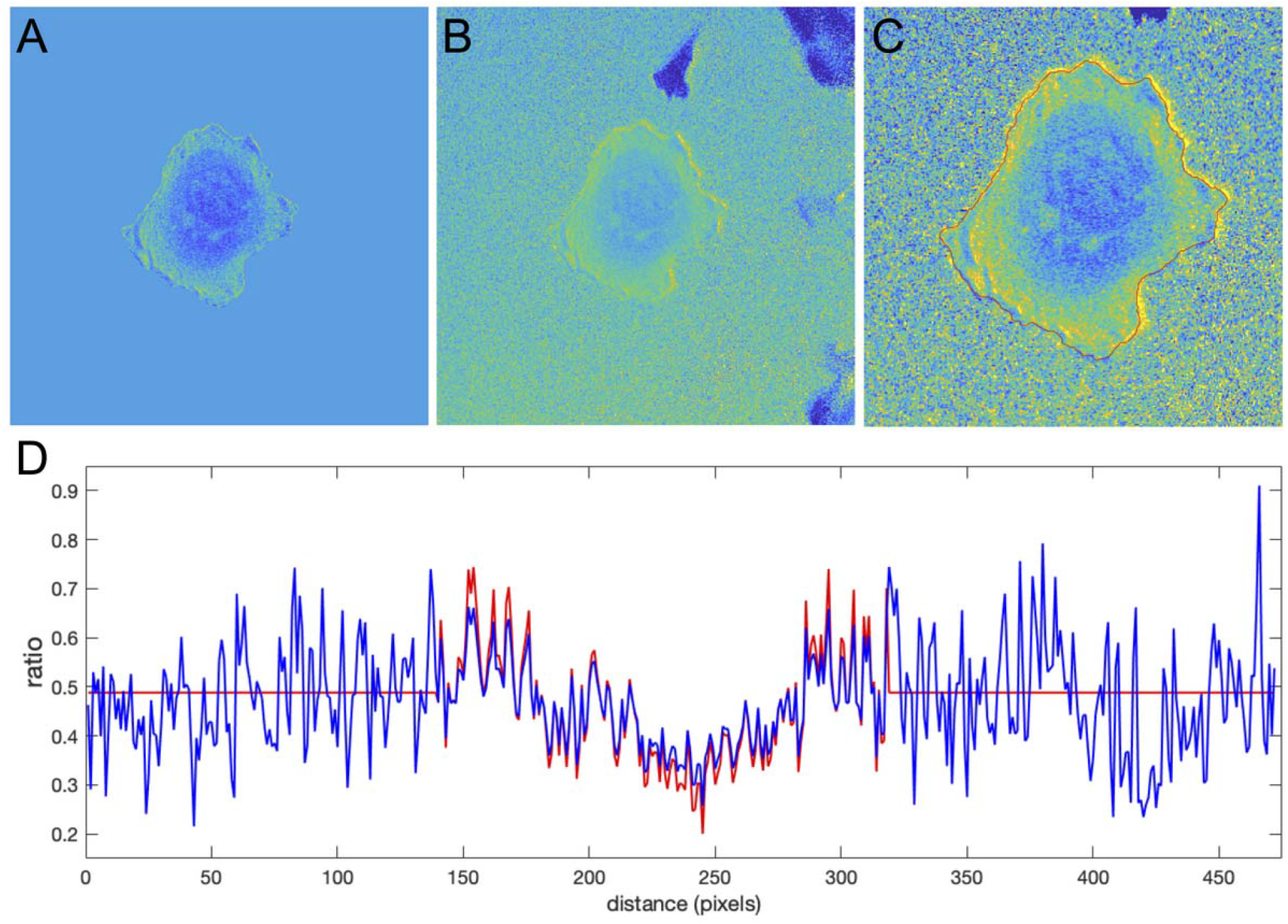
Application of the correction factor method to the Cdc42 biosensor data. **A**. The ratio image resulting from the mean background subtraction method. The background level is set to the mean value of the ratio inside the cell. **B**. The ratio image resulting from the correction factor method. Now with the ratio signal in the background, a shade-like bias from the bottom-left to the top-right sides of the cell becomes clearly visible. **C**. A zoomed-in image of the cell with the cell outline highlighted in red. The image shows that the artifact is mostly present outside the cell so that the ratio calculation inside the cell away from the edge should be accurate in both methods, but the bias is still present at the very edge. **D**. The ratio values along a horizontal line running across the whole image. Red and blue colors represent the results of the mean background subtraction and correction factor methods, respectively.

Despite the similarity of the intensity profiles between the methods, the correction method reveals that there is a ratio-imaging artifact. This artifact, the consistently higher ratio on one side of the cell is clearly visible near the cell edge. Now, after the fact, one may notice a hint of this bias in the ratio image from the MBS method, but only for a limited number of pixels at the very edge of the cell. Because the NCF method preserves the information near the edge outside the cell, the artifact is visible *much more distinctly*. This once again shows the practical benefits of using the NCF method. In summary, we found that CDC42 biosensor signal at the very edge of the cell (i.e., within 1 pixel from edge) is artificially biased and does not represent a biological process, so that the affected pixels should be excluded from further analysis. In published analyses, noise and resultant difficulties with ratio imaging at the very edge have led to exclusion of the outermost two pixels from analysis.

## Discussion

For biosensors, ratiometric imaging is used to exclude the effects of uneven illumination, nonhomogeneous biosensor distribution, etc. In this article, we show that this procedure can create noise-related artifacts at the cell edge where the fluorescence is weak and contains a substantial noise component. In the past, these artifacts have been eliminated or reduced by excluding pixels at the very edge from consideration, or by averaging larger regions of the cell edge (Pertz el at 2006, Machacek et al 2009), but these fixes sacrificed spatial information or resolution. The regions near the edge of protruding membranes are important from a biological, mechanistic perspective. We therefore provide here a simple approach to investigate the ratio at the very edge of the cell, to identify potential issues and even mitigate their effects. Importantly, this NCF method enables calculation of the ratio in the portion of the image outside the cell, because this no longer requires division by very low values after background subtraction. The ratio outside the cell can be used to determine an NCF value and thereby eliminate artifacts within the cell.

In general, the NCF method is based on a minimization routine that flattens the noise level in the ratio near the edge outside the cells. Alternatively, for cases when the background ratio in cells is significant even in the absence of protein activity (e.g., for biosensors with FRET in the off state), the NCF value can be found by minimizing the difference between NCF and MBS ratios in the cell region away from the edge. Using a mathematical analysis, we showed that both calculation methods give NCF ratio measurements that are consistent with traditional MBS approaches but free from the noise-related artifacts (because we eliminate the need to divide the FRET signal by small numbers). We validated the accuracy of the theoretical predictions using data from two uniformly distributed fluorophores. Our method allowed us to achieve the expected flat ratio with ∼1% accuracy, while the MBS method gave ∼5% deviation at best. In parts of the cell where the FRET signal is weak, the ratio values from the NCF method are leveled with the background noise, unlike the MBS method that does not give a direct reference point for zero activity.

When we applied our method to analyze the activity of the Asef biosensor, we discovered that there was persistent Asef activity in a very narrow band (2 pixel = 640 nm) on the cell edge. In contrast, deeper within the cell, GEF activity varied significantly during protrusion and retraction (data not shown). These intriguing spatial differences in GEF activity will be pursued in further studies. Knowing that the signal at the very edge of the cell is not an artifact due to calculating ratios at the limit of image resolution is a critical first step.

Applying our NCF method to the dual-chain Cdc42 biosensor revealed a different artifact in the time-lapse cell images, a spatially biased background signal near the cell edge. Although such uneven distribution affected mostly the pixels right outside the cell mask, the pixels on the edge of the mask were still impacted by this bias. This was not obvious when background regions of the image were “zeroed out” using the traditional MBS method, but was clearly apparent when ratios inside and outside the edge could be compared.

Importantly, identifying a correct NCF value not only removes artifacts stemming from low signal/noise, but also produces a flat level of background noise in the ratio image. This can potentially be useful for stabilizing drift in time-lapse recordings, such as that produced by photobleaching.

## Methods

### Live cell imaging

For the inactive biosensor experiments, Cerulean and YPet were transfected into Cos7 cells using Fugene6 (Roche) 24 hr prior to imaging. On the day of imaging, cells were trypsinized using Trypsin/EDTA (Corning). They were then replated onto coverslips coated with Fibronectin (10µg/ml 37C overnight) and allowed to attach in DMEM (Corning) /10%FBS (Hyclone). Cells were imaged in Hams/F12 (Caisson Labs)/5% FBS. Cells were imaged using a 40X, 1.3 NA objective on an Olympus IX-81 inverted microscope and using Metamorph screen acquisition software (Molecular Devices) and mercury arc lamp illumination. Filters used were Ex - ET436/20X, Em; donor- ET470/24M, FRET - ET535/30M and a 445/505/580 ET dichroic mirror. Images were obtained on a Flash4 sCMOS camera (Hamamatsu). Images were analyzed using MATLAB (Mathworks).

For the Asef and Cdc42 experiments, the Asef biosensor constructs were inserted into a tet-off inducible retroviral expression system and stable lines were produced in tet-off MDA-MB-231 cells (Johnson Lab, UNC-CH). Cells were maintained in DMEM (Cellgro) with 10% FBS (Hyclone) and 0.2 μg/ml doxycycline to repress biosensor expression. Biosensor expression was induced 48 hr prior to imaging through trypsinization and culturing without doxycycline and the Cdc42 biosensors were transfected into the RhoGEF biosensor-expressing stable cell lines. On the day of imaging, cells were replated using Accumax (Innovative Cell Technologies) onto coverslips coated with collagen I (10µg/ml 37C overnight) and allowed to attach in DMEM /10%FBS. After 2 hrs the media was replaced with Hams/F12 with 0.2% BSA, 10ng/µl Epidermal growth factor (R and D systems) 10mM HEPES, 100 µm Trolox, and 0.5mM Ascorbate and cells were allowed to equilibrate. After a further 2-4 hrs, cells were imaged in a closed chamber with media treated with Oxyfluor (1/100). For single biosensor experiments, cells were imaged using the filters listed above. For dual biosensor experiments, the excitation filters used were FF-434/17 for Cerulean3/mTFP and LSSmOrange, and FF-546/6 for Cherry (Semrock) combined with a custom zt440/545 dichroic (Chroma). For emission, a TuCam (Andor) was fitted with a FF560-FDi01 imaging-flat dichroic and a Gemini dual view (Hamamatsu) was added to each emission port. For the short wavelength port of the Gemini, the filters used were donor- FF-482/35, FRET - FF-520/15 and a FF509-FDi01 imaging flat dichroic mirror. For the Red-shifted Gemini port, the filters used were Orange – FF01-575/15, FRET/mCherry – FF01-647/57 and a FF580-FDi01 imaging flat dichroic.

### Image pre-processing

Donor and FRET images were aligned using fluorescent beads as fiduciaries to produce a transformation matrix using the Matlab function “cp2tform” (Matlab, The Mathworks Inc.). This was then applied to the Donor image using the Matlab function “imtransform”. The camera dark current was determined by obtaining images for each camera without excitation, and the dark current was subtracted from all images. Images were corrected for shading due to uneven illumination by taking images of a uniform dye solution under conditions used for each wavelength, normalizing this image to an average intensity of 1 to produce a reference image for each wavelength, and then dividing the images corrected for dark current by the shading correction reference image.

### NCF processing

MATLAB scripts for NCF application are available upon request

## Acknowledgements

This work was supported by grants from the U.S. Army Research Office (ARO, W911NF-17-1-0395) and the National Science Foundation (CMMI 1942561) to D.T, and by NIGMS grant GM-R35GM122596 to KMH.

## Supporting information

Supplemental Text

## References

Davies, E. R. (2005). Machine vision : theory, algorithms, practicalities. Amsterdam ; Boston, Elsevier.

Greenwald, E. C., S. Mehta and J. Zhang (2018). “Genetically Encoded Fluorescent Biosensors Illuminate the Spatiotemporal Regulation of Signaling Networks.” Chem Rev 118(24): 11707–11794.

Hall, A. (2012). “Rho family GTPases.” Biochem Soc Trans 40(6): 1378–1382.

Hochreiter, B., A. P. Garcia and J. A. Schmid (2015). “Fluorescent proteins as genetically encoded FRET biosensors in life sciences.” Sensors (Basel) 15(10): 26281–26314.

Hodgson, L., F. Shen and K. Hahn (2010). “Biosensors for characterizing the dynamics of rho family GTPases in living cells.” Curr Protoc Cell Biol Chapter 14: Unit 14 11 11-26.

Kurokawa, K., R. E. Itoh, H. Yoshizaki, Y. O. Nakamura and M. Matsuda (2004). “Coactivation of Rac1 and Cdc42 at lamellipodia and membrane ruffles induced by epidermal growth factor.” Mol Biol Cell 15(3): 1003–1010.

Machacek, M., L. Hodgson, C. Welch, H. Elliott, O. Pertz, P. Nalbant, A. Abell, G. L. Johnson, K. M. Hahn and G. Danuser (2009). “Coordination of Rho GTPase activities during cell protrusion.” Nature 461(7260): 99–103.

Marston, D. J., M. Vilela, J. Huh, J. Ren, M. L. Azoitei, G. Glekas, G. Danuser, J. Sondek and K. M. Hahn (2020). “Multiplexed GTPase and GEF biosensor imaging enables network connectivity analysis.” Nat Chem Biol 16(8): 826–833.

Pertz, O., L. Hodgson, R. L. Klemke and K. M. Hahn (2006). “Spatiotemporal dynamics of RhoA activity in migrating cells.” Nature 440(7087): 1069–1072.

Rossman, K. L., C.J. Der and J. Sondek (2005). “GEF means go: turning on Rho GTPases with guanine nucleotide-exchange factors.” Nat Rev Mol Cell Biol 6(2): 167–180.

Terai, K., A. Imanishi, C. Li and M. Matsuda (2019). “Two Decades of Genetically Encoded Biosensors Based on Forster Resonance Energy Transfer.” Cell Struct Funct 44(2): 153–169.

