## Supplemental Information for "Correcting artifacts in ratiometric biosensor imaging; an improved approach for dividing noisy signals"

**This PDF file includes:**

Supplemental Text

**Extended derivation of the mathematical expression in section "*A hypothetical cell model illustrating effects of noise on ratio values*":**

$$\frac{s_F x + b_F + n_F}{s_D x + b_D + n_D} = \frac{s_F\left(x + \frac{b_D}{s_D}\right) + \left(b_F - s_F\frac{b_D}{s_D}\right) + n_F}{s_D\left(x + \frac{b_D}{s_D}\right) + n_D}$$

Here, the first step is to add $\left(s_F\frac{b_D}{s_D} - s_F\frac{b_D}{s_D}\right)$, which is **zero**, to the numerator and multiply $b_D$ by $\frac{s_D}{s_D}$, which is **one**, in the denominator. Second step is combining terms. The last step is taking $s_F$ and $s_D$ out of brackets:

$$\frac{s_F x + b_F + n_F}{s_D x + b_D + n_D} = \frac{s_F x + \left(s_F\frac{b_D}{s_D} - s_F\frac{b_D}{s_D}\right) + b_F + n_F}{s_D x + \frac{s_D}{s_D}b_D + n_D} = \frac{s_F x + s_F\frac{b_D}{s_D} - s_F\frac{b_D}{s_D} + b_F + n_F}{s_D x + \frac{s_D}{s_D}b_D + n_D}$$

$$= \frac{\left(s_F x + s_F\frac{b_D}{s_D}\right) + \left(-s_F\frac{b_D}{s_D} + b_F\right) + n_F}{\left(s_D x + \frac{s_D}{s_D}b_D\right) + n_D} = \frac{s_F\left(x + \frac{b_D}{s_D}\right) + \left(b_F - s_F\frac{b_D}{s_D}\right) + n_F}{s_D\left(x + \frac{b_D}{s_D}\right) + n_D}$$

**Extended derivation of the mathematical expression in section "*Use of a noise correction factor; identification and correction of artifacts without using direct background subtraction*":**

$$Ratio(x,y) = \frac{image1(x,y)}{image2(x,y)} = \frac{a_0(S_2(x,y) + B_2) + (B_1 - a_0B_2) + N_1(x,y)}{(S_2(x,y) + B_2) + N_2(x,y)}$$

Here, the first step is to add $(a_0B_2 - a_0B_2)$, which is **zero**, to the numerator and combine the terms as highlighted in cyan and gray. The last step is taking $a_0$ out of brackets:

$$Ratio(x,y) = \frac{image1(x,y)}{image2(x,y)} = \frac{a_0S_2(x,y) + B_1 + N_1(x,y)}{S_2(x,y) + B_2 + N_2(x,y)}$$

$$= \frac{a_0S_2(x,y) + a_0B_2 - a_0B_2 + B_1 + N_1(x,y)}{S_2(x,y) + B_2 + N_2(x,y)}$$

$$= \frac{a_0S_2(x,y) + a_0B_2 - a_0B_2 + B_1 + N_1(x,y)}{S_2(x,y) + B_2 + N_2(x,y)}$$

$$= \frac{a_0(S_2(x,y) + B_2) + (B_1 - a_0B_2) + N_1(x,y)}{(S_2(x,y) + B_2) + N_2(x,y)}$$
